# Mismatch between pollen and pistil size causes asymmetric mechanical reproductive isolation across *Phlox* species

**DOI:** 10.1101/2024.05.08.593106

**Authors:** Anna F. Feller, Grace Burgin, Nia Lewis, Rohan Prabhu, Robin Hopkins

**Affiliations:** Department of Organismic and Evolutionary Biology & Arnold Arboretum, Harvard University, Cambridge, MA 02138, USA; Northeastern University, Boston, MA 02115, USA

**Keywords:** Phlox, reproductive isolation, pollen-pistil interactions, post-mating prezygotic barriers

## Abstract

In flowering plants, pollen-pistil interactions can serve as an important barrier to reproduction between species. As the last barrier to reproduction before fertilization, interactions between these reproductive organs are both complex and important for determining a suitable mate. Here, we test whether differences in style length generate a post-mating prezygotic mechanical barrier between five species of perennial *Phlox* wildflowers with geographically overlapping distributions. We perform controlled pairwise reciprocal crosses between three species with long styles and two species with short styles to assess crossing success (seed set). We find that heterospecific seed set is broadly reduced compared to conspecific cross success and reveal a striking asymmetry in heterospecific crosses between species with different style lengths. To determine the mechanism underlying this asymmetric reproductive isolating barrier we assess pollen tube growth *in vitro* and *in vivo*. We demonstrate that pollen tubes of short-styled species do not grow long enough to reach the ovaries of long-styled species. We find that short-styled species also have smaller pollen and that both within and between species pollen diameter is highly correlated with pollen tube length. Our results support the hypothesis that the small pollen of short-styled species lacks resources to grow pollen tubes long enough to access the ovaries of the long-styled species, resulting in an asymmetrical, mechanical barrier to reproduction. Such mechanisms, combined with additional pollen-pistil incompatibilities, may be particularly important for closely related species in geographic proximity that share pollinators.

## Introduction

The incredible diversity of floral traits –in size, shape, and color– is integral to diversification in angiosperms (Stebbins, 1970; Faegri & Van Der Pijl, 1979; Soltis & Soltis, 2014). The role of floral trait variation in the speciation process is particularly well-appreciated for its impact on pollination; variation in floral morphology is linked to variation in pollination vector or mode, which can decrease pollen flow between diverging lineages (Grant, 1949, 1994; Fenster *et al*., 2004; Kay *et al*., 2006; Waser & Ollerton, 2006; Van der Niet *et al*., 2014; Landis *et al*., 2018; Hernández-Hernández & Wiens, 2020; Stewart *et al*., 2022). Less widely explored is how variation in reproductive organs within the flower might generate mechanical barriers to gene flow during speciation. As overall flower size and / or shape diversifies, so does the size, shape, and placement of the male and female reproductive organs, which can drive mechanical mismatch during reproduction between species.

A mechanical mismatch in male and female reproductive organs in animals can cause reproductive isolation due to failed or suboptimal copulation (Garlovsky *et al*., 2023) and has been observed in a variety of taxa (e.g., in beetles (Soudi *et al*., 2016), millipedes (Tanabe & Sota, 2008), drosophila (Kamimura & Mitsumoto, 2012), damselflies (Sánchez-Guillén *et al*., 2012), mosquitofish (Anderson & Langerhans, 2015)). Mechanical mismatches in reproductive organs may be similarly important to speciation in plants. It can occur both during pollination as a mismatch in position of stigma and pollen deposition (Grant, 1994; Minnaar *et al*., 2019; Kay & Surget-Groba, 2022), as well as post-pollination through pollen-pistil interactions.

Following pollination, sexual reproduction in flowering plants requires complex and coordinated communication between pollen and pistil (reviewed in Bedinger *et al*. (2017); Zheng *et al*. (2018); Johnson *et al*. (2019); Robichaux & Wallace (2021); Cheung *et al*. (2022)). After arriving at the flower, pollen must germinate on the receptive stigma, grow a pollen tube through the style, locate ovules housed at the base of the pistil, and deliver sperm for fertilization. Throughout this post-pollination process, pollen receives molecular signals from receptive pistils to facilitate germination, initiate pollen tube growth, and guide pollen to the ovule (Zheng *et al*., 2018; Johnson *et al*., 2019; Robichaux & Wallace, 2021; Cheung *et al*., 2022; Zakharova *et al*., 2022). Species-specific biochemical signaling and active heterospecific rejection by the pistil tissue at any one of these post-pollination steps can cause reproductive isolation (Broz & Bedinger, 2021). In addition to biochemical signaling, mechanical isolating barriers at the pollen-pistil interface can also emerge as a result of divergence in floral traits between species, as for example in *Silene* (Nista *et al*., 2015).

One key floral trait is style length, which varies across flowering plants, sometimes significantly even between closely related species (e.g., Buchholz *et al*. (1935); Levin & Kerster (1967); Williams & Rouse (1988); Torres (2000); Aguilar *et al*. (2002); Nista *et al*. (2015)). By determining how far a pollen tube must grow to reach the ovules, differences in style length may serve as an important post-pollination mechanical isolating barrier (Tiffin *et al*., 2001). It has been observed that plants with long styles do not set successful seed when pollinated by plants with short styles while short-styled plants can successfully reproduce with pollen from long-styled plants. This asymmetry has been documented in a variety of closely related plant species (e.g., Perez & Moore (1985); Williams & Rouse (1988); Tiffin *et al*. (2001); Lee *et al*. (2008); Yost & Kay (2009); Nista *et al*. (2015)), demonstrating that variation in style length can be associated with a strong post-pollination mechanical isolating barrier across flowering plant diversity. However, we know little about how this barrier functions. Why are pollen grains from short-styled species unsuccessful at fertilizing ovules of long-styled species?

One hypothesis is that pollen from short-styled species may lack the resources to grow pollen tubes long enough to reach the ovules of long-styled species (Delpino, 1867; Nista *et al*., 2015; Brothers & Delph, 2017). Support for this hypothesis is largely correlative; longer-styled species tend to have larger pollen grains than shorter-styled species (e.g., Plitmann & Levin (1983); Roulston *et al*. (2000); Torres (2000); Aguilar *et al*. (2002)). If smaller pollen is less resourced, it could be unable to grow tubes long enough to reach ovules at the base of long styles. While there is broad acceptance for this hypothesis, it has rarely been demonstrated conclusively (but see Nista *et al*. (2015) and Brothers & Delph (2017)). An alternative hypothesis is that the pistil environment of short-versus long-styled species varies, generating a mismatch in biochemical signaling between pollen from short-styled species and long-styled pistils (Amici, 1830; Cruden & Lyon, 1985; Swanson *et al*., 2004; Cruden, 2009; Broz & Bedinger, 2021; Kato *et al*., 2022). Distinguishing between these two scenarios requires disentangling the effect of pollen size/resources from variation in the pistil environment across species.

Here, we investigate the mechanisms driving mechanical isolation between long and short styled species of *Phlox* wildflowers. *Phlox* is a genus of flowering plants with more than 60 species that are predominantly found in Northern America (Wherry, 1955; Ferguson & Jansen, 2002). The species used in this study, *P. glaberrima* ssp. interior (GLA), *P. maculata* (MAC), *P. paniculata* (PAN), *P. pilosa* spp. pilosa North (PIL), *P. divaricata* (DIV; including the two subspecies *P. divaricata* ssp. divaricata and *P. divaricata ssp*. laphamii) are all perennial species that evolved in the prairies of the midwestern United States, still occur with widely overlapping geographic distributions in the states of Illinois and Indiana, and likely share pollinators (Robertson, 1891, 1895, 1928; Wherry, 1932, 1933, 1955) (Figure 1). Three of the species (GLA, MAC, PAN) have long styles that protrude from the calyx and two (DIV, PIL) have short styles that are hidden deep within the calyx.

**Figure 1.**
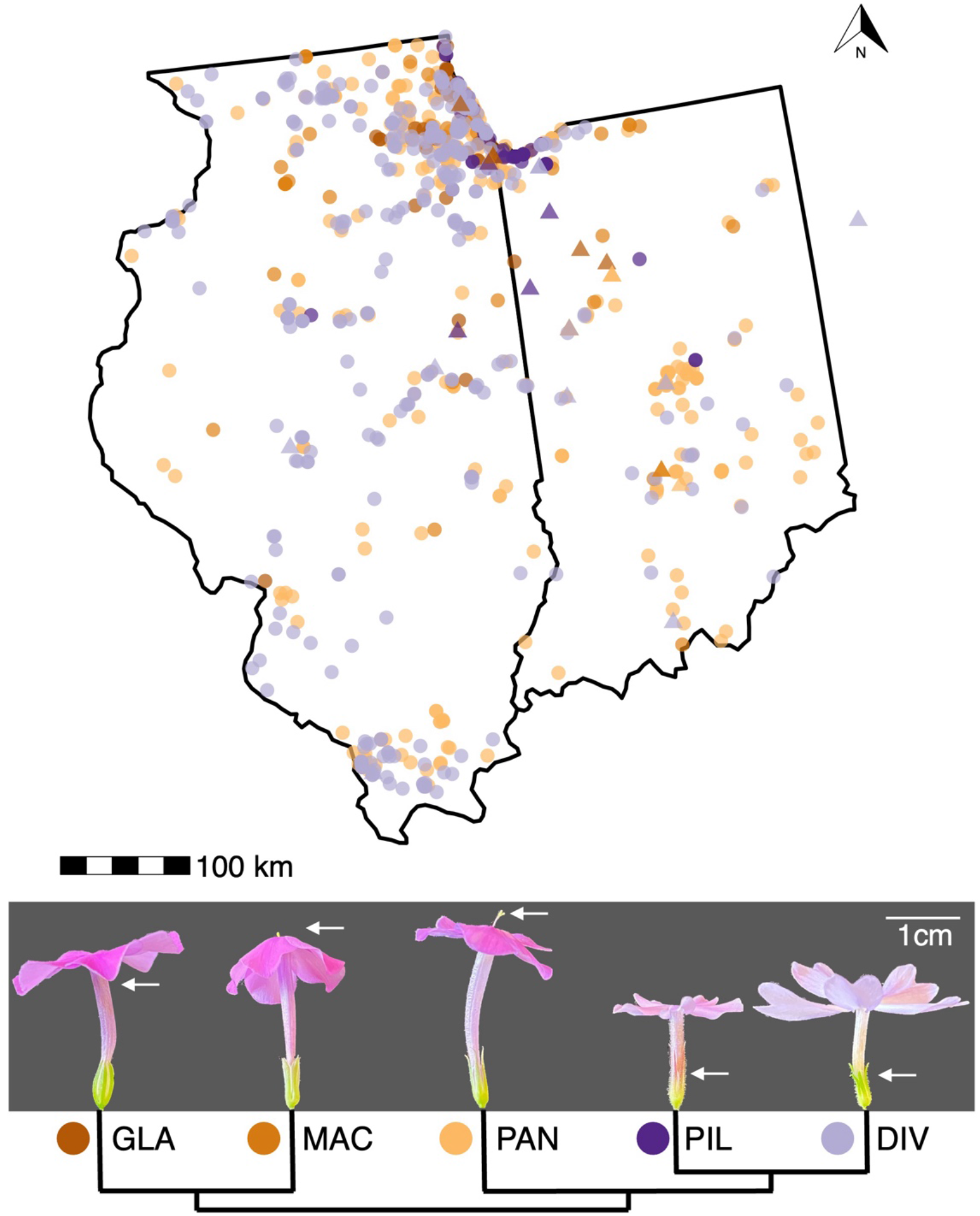
Distributions of the five focal *Phlox* species with geographically overlapping ranges in the states of Indiana and Illinois (USA). Occurrence data was obtained from (GBIF.org, 2023; see Table S1 for the full reference list). Triangles indicate the locations of individuals used in this study (including one DIV in Ohio; see Table S2). The map was plotted with the R packages usmap v.0.6.1 (Di Lorenzo, 2023), ggplot2 v.3.4.2 (Wickham, 2016), and ggspatial v.1.1.9 (Dunnington, 2023). *P. glaberrima* ssp. interior (GLA), *P. maculata* (MAC), and *P. paniculata* (PAN) are long-styled. *P. pilosa* spp. pilosa North (PIL), and *P. divaricata* (DIV) are short-styled. The white arrows indicate the approximate height of styles for each species.

We first establish that style length variation is a post-mating prezygotic barrier between short- and long-styled species by performing pairwise reciprocal crosses to assess crossing success (seed set). Second, we measure a suite of floral morphological traits to determine the extent of floral divergence associated with style length variation. Finally, we investigate the mechanism of fertilization failure by assessing pollen tube growth *in vivo* on cross-pollinated pistils as well as *in vitro* on a standardized medium.

## Methods

### Study plants

Plants were collected from natural populations during the summers of 2017-2019 and 2022 (Table S2). We propagated vegetative clones from cuttings of field-collected plants under controlled conditions in greenhouses at 23 degrees Celsius during the day, 20 degrees Celsius at night, and 50% relative humidity. When needed, plants were vernalized for six weeks at 7.2 degrees Celsius to induce flowering.

### Crossing experiment for seed set

To assess the degree of successful sexual reproduction between and within species, we quantified seed set from controlled cross-pollinations. We performed all pairwise reciprocal crosses between our five focal species, including conspecific crosses, with three replicate individuals of each pair in each direction for a total of 75 cross combinations. For each cross combination, we pollinated 10 replicate flowers (except for nine cross combinations where only 4-9 flowers were available to cross; see Table S3). Conspecific crosses were performed between individuals from different populations. To prevent self-pollination, we removed petals and the attached anthers from buds that were predicted to open within 24-48 hours. Three days after anther removal, it was confirmed that stigmas were receptive (all three lobes clearly visible and separated) before transferring pollen onto them using forceps. We used light mesh fabric to bag pollinated flowers so we could collect and count seeds that would otherwise be explosively dehisced. We counted seeds produced from fully ripened fruits.

We used R v.4.2.3 (R Core Team, 2023) to visualize and analyze the results. Functions in the R packages dplyr v.1.1.2 (Wickham *et al*., 2023) and here v.1.0.1 (Müller, 2020) were used for data processing. We quantified seed set as the average number of seeds per flower obtained across three replicates of each species combination and direction of cross. We calculated RI following (Sobel & Chen, 2014) using their Equation 4A:

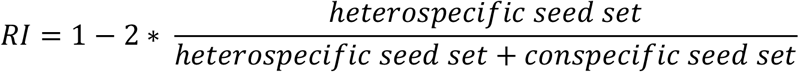

### Floral trait measurements

We used five to nine mature flowers of three to five individuals per species to measure style length, calyx height, tube length, petal limb length, and corolla area (see Figure 3). The flowers were adhered to white paper with book tape and 2D-scanned using an EPSON Perfection V600 Photo scanner. A scale and color bar were included in each scan. For the style length and calyx height measurements, the anthers and petals were removed from the flowers. We measured all traits in ImageJ 1.53k (Rasband, 1997-2018) after setting the scale using the straight line tool, or the color thresholding and analyze particles functions for the corolla area, respectively. To statistically assess the differences between long- and short-styled species, we built mixed effect models with ‘type’ (style length long vs short) as fixed effect and ‘individual’ nested within ‘species’ as random effect. We used the R packages lme4 v. 1.1-34 (Bates *et al*., 2015) to fit the models. To test the significance of the fixed effects, we compared the model with and without the fixed effect using the anova function (with REML=F).

### Pollen size quantification

We quantified pollen size of three individuals per species, and three flowers per individual. We transferred the mature anthers of a single flower into a 1.5 mL Eppendorf tube containing 50 μl of 1x phosphate-buffered saline (PBS). We vortexed the tube for 10 seconds to liberate the pollen from the anthers and transferred 10 μl of the PBS with pollen into one chamber of a 2-Chip hemocytometer by Bulldog Bio. We used a ZEISS Axio Imager.A2 microscope with ZEN 2.3 pro software to take standardized photos with a 10x objective. We batch-processed the resulting TIFF images in ImageJ 1.53k (Rasband, 1997-2018) using a customized macro based on (“Building an imagej macro for batch processing of images from imaging well plates,” 2019). In brief, the images were converted to grayscale, thresholded using default parameters, and then the ‘analyze particles’ function was used to quantify the pollen area. The scale was set globally on one image before running the macro. The results were statistically analyzed using the same structure of mixed-effect models as described for the floral trait measurements.

### In-vitro pollen tube quantification

To compare the growth of pollen from short- and long-styled species in a neutral environment, we germinated pollen in an artificial growth medium (Rodriguez-Enriquez *et al*., 2013). We used the following concentrations: 10% sucrose, 0.1 mM spermidine, 10 mM GABA, with an adjusted pH of 7.5. We divided microscope slides (Fisher Scientific) into three compartments using an ImmEdge® Hydrophobic Barrier PAP Pen (H-4000) and filled each with 125 μl of medium. (We did not use cellulose membranes.) Prepared slides were used on the same or next day, following the storage and acclimation procedures in Rodriguez-Enriquez *et al*. (2013). We used fine tip (size #00) paint brushes to apply pollen to the medium. The slides were then incubated in petri dishes with a wet tissue for 24 hours at 24 degrees Celsius in the dark. After incubation we transferred them to 4 degrees Celsius until imaging (which we performed immediately). We used a ZEISS Axio Imager.A2 microscope with ZEN 2.3 pro software for imaging. To be quantified, the pollen tubes had to fulfill three criteria: 1) be at least three times the length of the pollen grain; 2) the beginning and end had to be clearly visible; 3) the whole tube length had to be mostly in the same z-plane.

From each pollen grain that passed our criteria we took paired measurements of pollen diameter and pollen tube length. We used the straight-line tool in ImageJ to measure pollen diameter and the segmented line tool to measure the pollen tubes. The results were visualized and statistically analyzed using similar mixed models as above with style length as fixed effect and individual nested within species as random effect. Additionally, we performed correlation tests between pollen diameter and pollen tube length by species and across all species.

### In-vivo pollen tube quantification

We used the same crossing procedure as described above in the seed set experiment to obtain images of hetero- and conspecific pollen tubes growing in pistils. We experimentally verified that all conspecific pollen tubes were able to reach conspecific ovules in 48 hours. The crossed flowers (style and calyx) were removed from the plants after 48 hours, transferred into glass scintillation vials, and covered with a formalin-aceto-alcohol (FAA). After 24 hours, the FAA was exchanged with 70% ethanol and the vials were stored at 4 degrees Celsius until further processing. After a minimum of 24 hours, the ethanol was removed and 8M sodium hydroxide added. After 24 hours in sodium hydroxide, we dissected out the pistils from the calyces and stained them in 0.1% Aniline Blue for three hours. The pistils were then mounted on microscope slides in 50% glycerol. Pistils were visualized in the DAPI channel (358 nm excitation) using a ZEISS Axio Imager.A2 microscope with ZEN 2.3 pro software. Pistils were imaged in either 5x or 10x magnification, depending on style length. For long-styled species, multiple fields-of-view were stitched together using the ZEN 2.3 pro software.

## Results

### Seed set is asymmetric

Our results confirm that long-styled plants will not set seed with pollen from short-styled species. We found that seed set was generally lower in all heterospecific crosses compared to conspecific crosses (Figure 2, Tables 1 and S4). In all five species, a single flower can set a maximum of three seeds. The average number of seeds in conspecific crosses ranged from 1-1.5 seeds per flower and was <1 in all heterospecific crosses. In heterospecific crosses between different long-styled species (long x long) or between the two short-styled species (short x short), some seeds were obtained in both directions (with the exception of MACxPAN; Figure 2). In heterospecific crosses between species with different style lengths, seeds developed only in crosses when pollen from long-styled species was applied to a short-styled species (short x long). When pollen from short-styled species was applied to long-styled species (long x short), no seeds were produced (Figure 2, Table S4). This pattern of asymmetry is reflected in the RI values (Table 1). RI equals 1 (fully reproductively isolated) in all long x short crosses, and values below 0.5 are exclusively found in either long x long heterospecific crosses or short x long crosses.

**Table 1.** Post-pollination reproductive isolation values for each heterospecific cross with the first four crosses between species with similar style lengths, and the remaining (shaded) between species with different style lengths. (See also Table S4.)

| Species A | Species B | RI when pollen recipient is species A | RI when pollen recipient is species B |
| --- | --- | --- | --- |
| DIV (short) | PIL (short) | 0.670 | 0.909 |
| GLA (long) | PAN (long) | 0.686 | 0.415 |
| GLA (long) | MAC (long) | 0.365 | 0.262 |
| MAC (long) | PAN (long) | 1.000 | 0.871 |
| DIV (short) | GLA (long) | 0.368 | 1.000 |
| DIV (short) | MAC (long) | 0.916 | 1.000 |
| DIV (short) | PAN (long) | 0.876 | 1.000 |
| PIL (short) | GLA (long) | 0.377 | 1.000 |
| PIL (short) | PAN (long) | 0.750 | 1.000 |
| PIL (short) | MAC (long) | 1.000 | 1.000 |

DIV, *P. divaricata* (including subspecies *divaricata* and *laphamii*); GLA *P. glaberrima* spp. Interior; MAC, *P. maculata*; PAN, *P. paniculata*; PIL, *P. pilosa* spp. *pilosa*.

**Figure 2.**
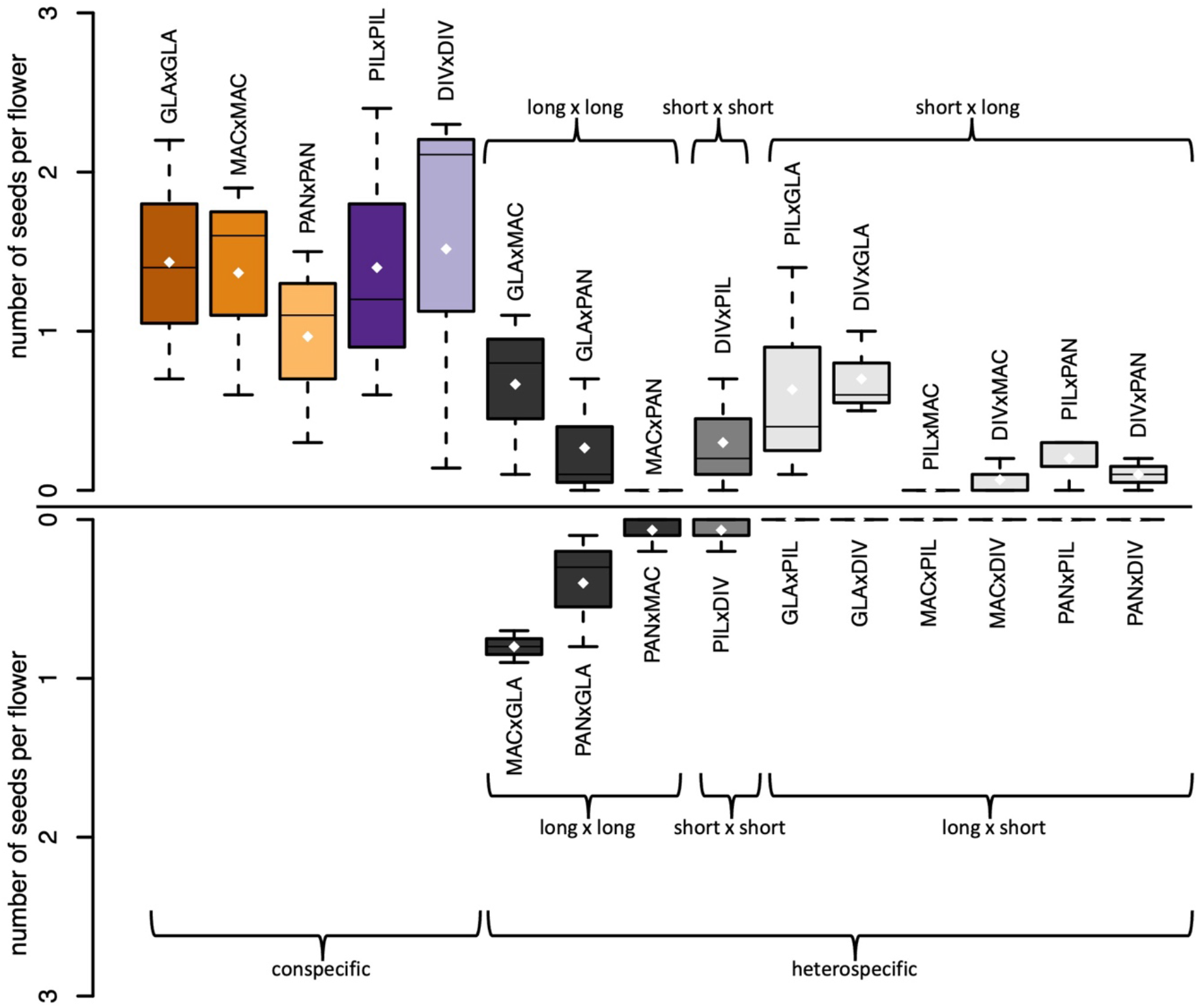
Seed set is reduced in heterospecific vs conspecific crosses and asymmetric in crosses between long- and short-styled species. Each cross consists of three replicates with up to ten crossed flowers per replicate (see Table S3). Shown is the average number of seeds obtained per flower in each replicate. The maximum number of seeds per flower is three. The white dots indicate the means. Species abbreviations are given as pollen-receiving species x pollen-donor species.

### Pollen size and some floral traits vary with style length

Style length (χ^2^(1) = 13.09, P = 0.0003), corolla tube length (χ^2^(1) = 4.21, P = 0.04), pollen diameter (χ^2^(1) = 12.23, P = 0.0005), and corolla area (χ^2^(1) = 6.4346, P = 0.011) all differed significantly by style-type (Table 2, Figure 3). Calyx height (χ^2^(1) = 0.54, P = 0.463), and petal limb length (χ^2^(1) = 0.04, P = 0.840) did not differ significantly by style-type (Table 2, Figure 3).

**Figure 3.**
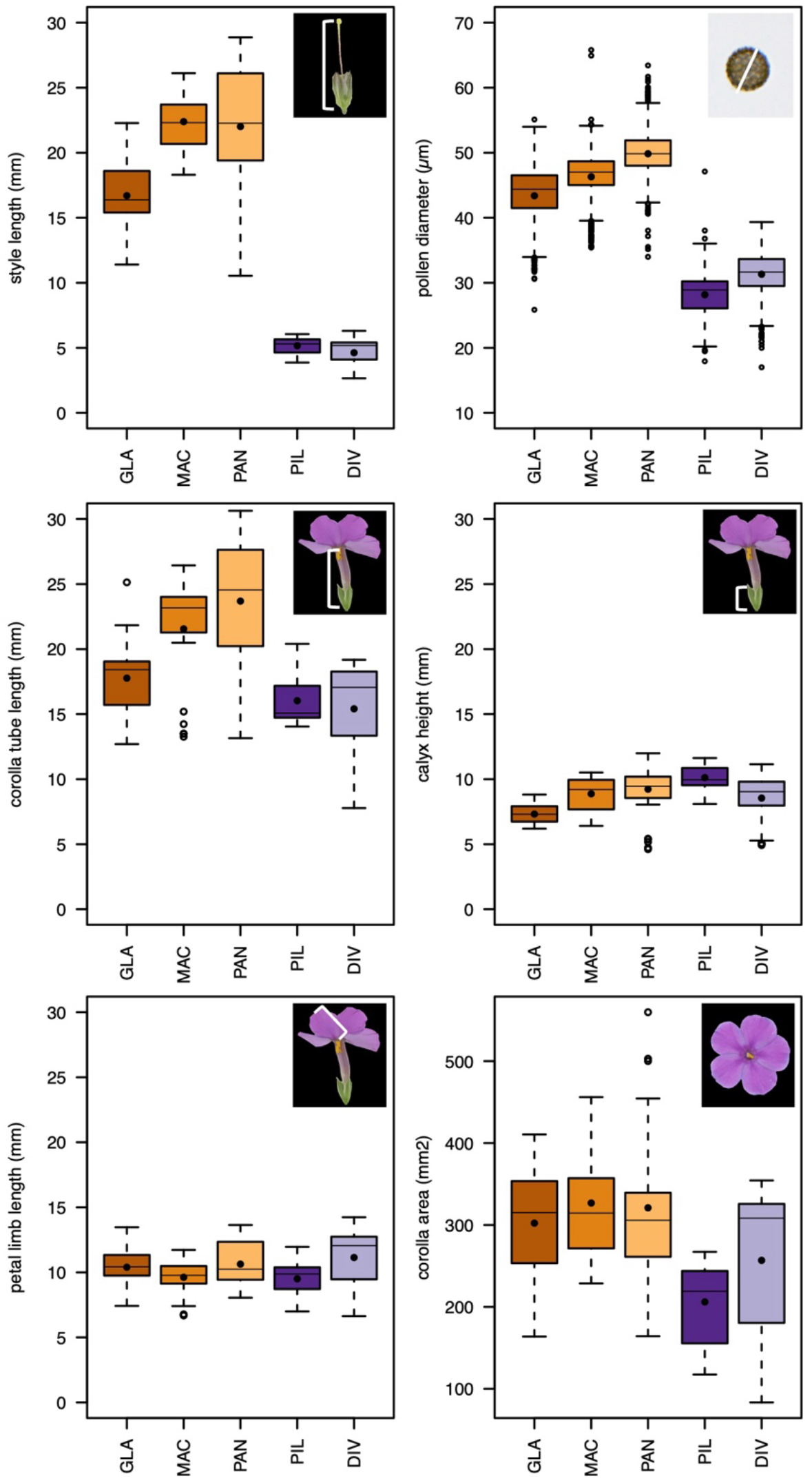
Long-styled species have larger pollen than short-styled species while other floral traits are similar. Five to nine flowers on three to five individuals per species were measured for the flower morphological traits, and pollen grains of three flowers of three individuals per species. The black dots indicate the means.

**Table 2.** Results from mixed effect models of. the continuous floral trait measurements with individual nested within species as random effect.

| Trait | Fixed effect | Estimate | ± | SE | df | $\chi^2$ - value | P-value |
| --- | --- | --- | --- | --- | --- | --- | --- |
| style length | Intercept (long style) | 20.31 | ± | 1.57 |  |  |  |
|  | short style | -15.51 | ± | 2.55 | 1 | 13.09 | 0.0003 *** |
| calyx height | Intercept (long style) | 8.47 | ± | 0.67 |  |  |  |
|  | short style | 0.65 | ± | 1.11 | 1 | 0.54 | 0.4630 |
| corolla tube length | Intercept (long style) | 20.74 | ± | 1.59 |  |  |  |
|  | short style | -5.38 | ± | 2.66 | 1 | 4.21 | 0.0401 * |
| petal limb length | Intercept (long style) | 10.25 | ± | 0.39 |  |  |  |
|  | short style | -0.13 | ± | 0.69 | 1 | 0.04 | 0.8419 |
| corolla area | Intercept (long style) | 314.23 | ± | 19.95 |  |  |  |
|  | short style | -91.36 | ± | 34.60 | 1 | 6.43 | 0.0112 * |
| pollen diameter | Intercept (long style) | 46.29 | ± | 1.84 |  |  |  |
|  | short style | -16.39 | ± | 2.91 | 1 | 12.23 | 0.0005 *** |
| pollen tube length | Intercept (long style) | 667.04 | ± | 54.94 |  |  |  |
|  | short style | -281.60 | ± | 87.39 | 1 | 7.38 | 0.0066 ** |

Significance codes: \*\*\*<0.001, \*\*<0.01, \*<0.05, .<0.1

### In-vitro pollen tube length correlates with style size and pollen size

Pollen tube length grown in-vitro for 24 hours on a standardized medium differed significantly by style-type (χ^2^(1) = 7.38, P = 0.007; Table 2, Figure 4). The length of the pollen tubes was highly correlated with pollen diameter across all species (r(643) = 0.429, P < 2.2e-16) as well as within species (GLA: r(129) = 0.235, P = 0.007; MAC: r(116) = 0.207, P = 0.024; PAN: r(97) = 0.286, P = 0.004; PIL: r(186) = 0.269, P = 0.0002; DIV: r(107) = 0.279, P = 0.003).

**Figure 4.**
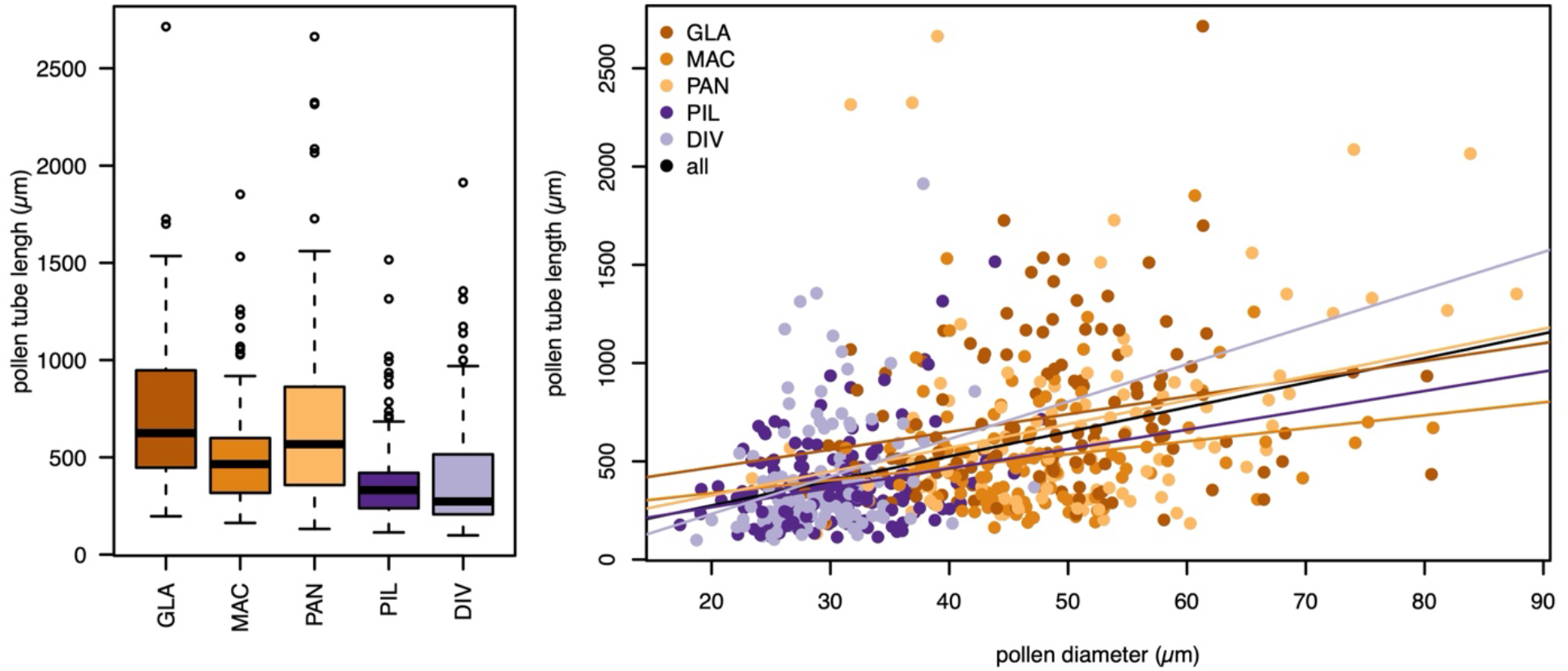
Pollen from long-styled species grow longer tubes *in vitro* and pollen tube length is significantly correlated with pollen grain size within and across species. Pollen grains were germinated and allowed to grow pollen tubes for 24 hours in a standardized medium. 13 to 98 pollen grains and tubes were measured of two to three individuals per species.

### In-vivo pollen tube growth is asymmetric

In crossed styles, we observed some pollen tubes from long-styled species reaching and entering the ovaries in short-styled species, even overshooting (Figure 5a). Pollen tubes from short-styled species did not reach the ovaries in long-styled species (Figure 5b).

**Figure 5.**
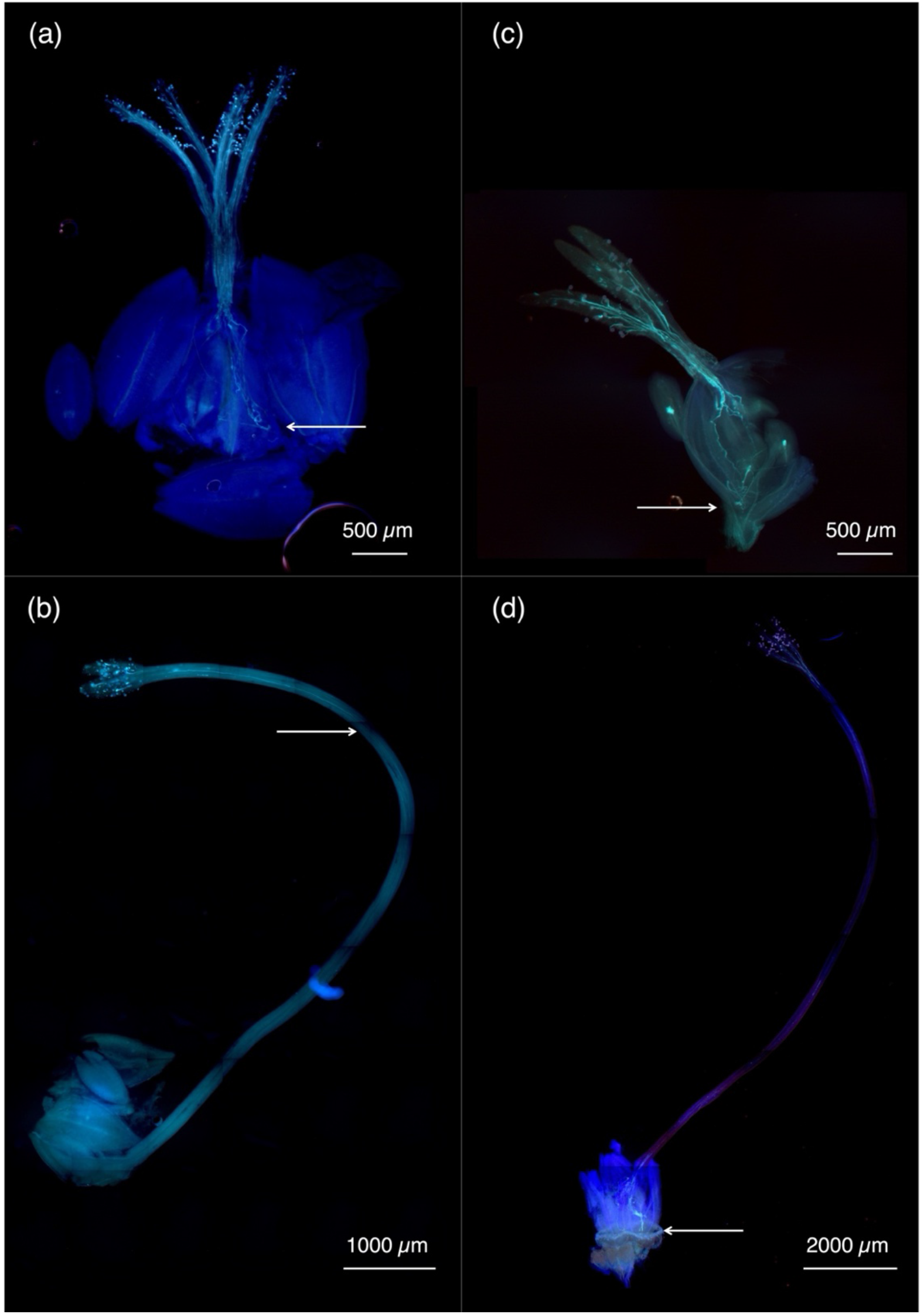
Representative images of pollen tubes grown in vivo. **a)** A representative image of a short style conspecific cross. **b)** Pollen tubes of short-styled species cannot grow into the ovaries of long-styled species, whereas **c)** pollen tubes of long-styled species can grow into the ovaries of short-styled species, even overshooting. **d)** A representative image of a long style conspecific crosses. The arrows indicate how far the pollen tubes grew within 48 hours.

## Discussion

We used a combination of crossing experiments, pollen tube growth assays, and morphological measurements to document the importance of post-mating prezygotic mechanical isolation in five species of *Phlox* wildflowers with varying style lengths. By growing pollen tubes in a standardized medium, our study controlled for variation in the pistil environment to further reveal resource properties of pollen as the underlying mechanism driving asymmetric crossing success between species of different style length.

The crossing results presented here corroborate those of previous experiments involving these *Phlox* species (Levin, 1963, 1966; Levin & Smith, 1965; Zale, 2014): heterospecific crosses yielded fewer seeds than conspecific crosses, and crosses between species with different style lengths yielded seeds only when the pollen parent was long-styled. As in these previous experiments, we found that controlled crosses within a species often did not result in complete seed set. This phenomenon could be because greenhouse conditions are not optimal for fertilization or seed maturation, or a normal developmental constraint that flowers tend to only mature seeds in a subset of available ovules. Unfortunately, no data exists on average seed set per flower in natural populations to compare to our controlled crosses in the greenhouse. However, central to our study is the relative comparison of conversus heterospecific seed set. Because all crosses were conducted under a common-garden environment, we expect no systematic bias between cross types.

Asymmetric crossing success between species with divergent style length has been documented in several systems (Perez & Moore, 1985; Williams & Rouse, 1988; Lee *et al*., 2008; Yost & Kay, 2009; Nista *et al*., 2015). In these systems, as in *Phlox*, pollen from short-styled species never fully reaches the ovaries of long-styled species following controlled pollination, whereas the opposite cross permits pollen success. The underlying mechanism driving this reproductive isolating barrier is difficult to disentangle using controlled pollinations alone. The failure of short pollen on long pistils could be caused by a lack of resources in the pollen (Delpino, 1867; Nista *et al*., 2015; Brothers & Delph, 2017), or by biochemical interactions in the pistil environment (e.g., spatial distribution of signaling molecules) (Broz & Bedinger, 2021). To disentangle the biochemical effect of the pistil environment from resource properties of the pollen grains, we grew pollen tubes in a standardized agar medium, thereby allowing pollen tubes to grow without the influence of a particular pistil environment. First, we found that although there is considerable variation within species, pollen from short-styled plants is significantly smaller than pollen from long-styled plants. We then found that when allowed to grow for the same amount of time, pollen from short-styled species grow significantly shorter pollen tubes than pollen from long-styled species. Finally, we found that pollen size and pollen tube length are correlated not only across species but also within each of the species we examined; smaller pollen makes smaller tubes both within and between species. Our data support the conclusion that pollen tubes from short-styled plants are limited by available resources in the small pollen-grains and thus cannot successfully reproduce as pollen-donors to long-styled plants.

*Phlox* is a particularly powerful system to investigate the mechanism underlying style-length reproductive isolation because flower morphology across this genus (and particularly in the five species investigated here) is broadly conserved. Unlike in other systems (e.g., Brothers & Delph (2017); B. Lanuza *et al*. (2023)), style length across our species is not associated with an overall difference in flower size. As a result, our study isolates the specific effect of style length variation as a reproductive isolating barrier from other floral trait changes. Although we do see minimal shifts in corolla tube length, the drastic difference between long and short styled *Phlox* is associated with a change in position of the stigma relative to the corolla tube opening but not a change in the overall shape and size of the flower (Figures 1 and 3). Based on phenotypic variation, it appears that the evolution of style length and pollen size are correlated with each other, but independent of variation in size of petals, calyx, and even stamen. Our findings of the partial decoupling of correlated evolution across floral organs motivates future work to determine if this is due to patterns of differential pleiotropy in the genetic basis of size variation across these floral organs, or due to patterns of selection acting differentially or in consort on different organs. Understanding why the pollen and style size seem to diverge together would provide an even greater understanding of why and how these mechanical barriers to reproduction evolved.

One of the primary explanations for why style length variation is associated with a shift in floral size is through the evolution of selfing syndrome traits following mating system transitions. Selfing species typically have an overall reduction in floral size relative to closely related outcrossing species (Sicard & Lenhard, 2011; Cutter, 2019; B. Lanuza *et al*., 2023). Importantly, these evolutionary correlations complicate interpretations of the specific underlying mechanism driving reproductive isolation. Transitions to self-fertilization are also associated with patterns of asymmetric post-mating prezygotic reproductive isolation due to biochemical mechanisms. A classic asymmetric crossing barrier in flowering plants is caused by an overlap in self- and heterospecific pollen rejection mediated by shared molecular pathways (Hancock *et al*., 2003; Li & Chetelat, 2015; Roda & Hopkins, 2019). This overlap in pollen rejection has manifested a pattern of self-incompatible species (often with longer styles) being able to reject pollen from self-compatible species but not vice versa. In *Phlox* however, both the long- and the short-styled species are self-incompatible, maintain similarly showy flowers to attract pollinators, and are predominantly outcrossing (Levin, 1963, 1966). The conservation of mating system across these five species further isolates the role of style length variation in reproductive isolation. Because all species in our study express an active self-incompatibility mechanism, we expect that all species are also equally able to deploy heterospecific-incompatibility through pollen recognition and rejection mechanisms. We find additional support for this expectation from our controlled crossing data. Crossing success was reduced in all heterospecific crosses, even those between species of similar style lengths. Pollen from long-styled species only sometimes resulted in seed set when used to pollinate a short-styled species, indicating the presence of additional post-mating prezygotic reproductive isolating mechanisms. The mechanical barrier driven by style length differences in *Phlox* likely acts in concert with these other post-mating prezygotic isolating barriers. We observed low germination rates of pollen on heterospecific stigmas, and germinated pollen on heterospecific stigmas often did not result in pollen tube growth. This finding is not surprising considering that successful fertilization requires a suite of sequential interactions between pollen and pistil, each of which can act as a barrier if the previous fails (reviewed in Broz & Bedinger (2021)). These pollen-pistil barriers may be important in preventing gene flow across *Phlox* species broadly, while style length variation prevents interspecific gene flow in just one crossing direction.

Style-length evolves as a barrier to reproduction in the context of selection on traits affecting other possible prezygotic reproductive barriers. As discussed above, a shift in mating system from outcrossing to selfing can result in the evolution of correlated suites of floral traits including shrinking flower size and thus style length. In these cases, the mechanical barrier caused by style length variation is a side-effect of other major shifts in reproductive strategy and acts in concert to cause reproductive isolation. Similarly, style length may diverge as floral form evolves to accommodate a new type of pollinator (e.g., a shift from bee to hummingbird pollination) (Grant, 1994; Fenster *et al*., 2004; Waser & Ollerton, 2006; Yost & Kay, 2009; Hermann *et al*., 2015). Major changes in petal morphology and selection for pollen placement on pollinators can result in the evolution of style length variation. In this case, style length differences act together with a change in pollinator vector to cause reproductive isolation. Interestingly, in these *Phlox* species there is little evidence for pollinator-mediated barriers to reproduction. All five species in our study have broadly geographically overlapping ranges (Wherry, 1955; Figure 1), and some even grow in immediate vicinity to each other. They have distinct flowering time peaks but broadly overlapping flowering periods, ‘distinct but not mutually exclusive habitat requirements’ (Hadley & Levin, 1969), and likely the same pollinators (Robertson, 1891, 1895, 1928; Wherry, 1932, 1933; Grant & Grant, 1965). Pollen exchange has been documented in detail between GLA and PIL (Levin & Smith, 1965; Levin & Kerster, 1967) and likely occurs between the other species as well. The mechanical inability of pollen of short-styled species to fertilize long-styled species does not seem to be the side-effect of correlated shift in reproductive strategy (e.g., pollination or mating system), but clearly provides a strong, asymmetric, and likely important mechanical prezygotic barrier to gene exchange between these species in the wild.

Because the variation in style length across species is not associated with other major transitions in reproductive strategy, it is challenging to infer what drove the evolution of style length variation in this system. Across the phylogeny of *Phlox* most species have long-styles, and the monophyletic clade of short-styled species is nested within the broader group of long-styled species (Garner *et al*., 2024). This phylogenetic pattern suggests that the ancestor of the short-styled *Phlox* had a long style, and that short-style/small pollen is a derived state having evolved just once. Therefore, as style length was evolving from long to short, individuals with short styles would have been unable to pollinate long-styled plants but would still be accessible to (and potentially out-competed by) pollen from long-styled individuals (McCallum & Chang, 2016). This scenario would present a likely reproductive disadvantage to short-styled plants relative to long-styled individuals. We therefore hypothesize that style length evolved due to selection for some other selective purpose and not through selection for enhanced reproductive isolation. As selection drove the evolution of short styles/small pollen, reproductive isolation emerged as a byproduct. Future studies are necessary to better understand what might have driven the evolution of shorter styles.

Asymmetries in reproductive isolation in plants are most commonly observed at the postzygotic stage or at the post-pollination prezygotic stage (reviewed in Christie *et al*. (2022) and Tiffin *et al*. (2001)). At the post-pollination prezygotic stage, reproductive isolation is usually due to differences in mating systems or style lengths (Christie *et al*., 2022). We have shown that the pattern of asymmetry in the *Phlox* system is caused by a mechanical mismatch between pollen size and style length driven by the inability for less-resourced pollen to grow tubes long enough to traverse long-styled pistils. Such asymmetries affect the direction of gene flow and can impact introgression dynamics (Tiffin *et al*., 2001; Christie *et al*., 2022). Next steps should thus determine if these asymmetries translate into asymmetric patterns of gene flow between species. In general, prezygotic reproductive isolation tends to be stronger than postzygotic reproductive isolation in plants (Lowry *et al*., 2008; Christie *et al*., 2022), suggesting that the impact of pollen-pistil incompatibilities is likely important for determining patterns of gene flow across species boundaries.

## Data accessibility

Data and R-scripts will be available from the Dryad digital repository under https://doi.org/10.5061/dryad.fttdz091s upon acceptance of the manuscript.

## Author contributions

Conceptualization: AFF, RH; Methodology: AFF, GB, NL, RH; Formal analysis: AFF; Investigation: AFF, GB, NL, RP; Resources: RH; Data Curation: AFF; Writing - Original Draft: AFF, GB, RH; Writing - Review & Editing: AFF, GB, NL, RP, RH; Visualization: AFF, GB, NL; Supervision: AFF, RH; Project administration: AFF, RH; Funding acquisition: AFF, RH.

## Acknowledgments

Many thanks to Austin Garner for providing his insights on the Midwestern perennial Phlox; to Izzy Acevedo for initiating the propagation of the experimental plants; to Megan Ardolino, Lee Toomey, Scott Pedemonte, and Mike Barrett for their work in the greenhouses; and to the Hopkins lab group for many helpful discussions and comments. We thank the members of the following departments and parks for issuing permits for the 2022 sample collection to AFF and for facilitating access: Indiana Department of Natural Resources, Eagle Creek Park (IN), Illinois Department of Natural Resources, Illinois Nature Preserves Commission, Champaign County Forest Preserve District (IL), Grand Prairie Friends (IL), Forest Preserves of Cook County (IL), Johnny Appleseed Metropolitan Park District (OH).

