## Supplementary Materials for "Mismatch between pollen and pistil size causes asymmetric mechanical reproductive isolation across *Phlox* species"

**Table S1. GBIF references and download links**

| Species | References |
|---|---|
| *P. divaricata* ssp. divaricata (DIV) | iNaturalist contributors, iNaturalist (2022). iNaturalist Research-grade Observations. iNaturalist.org. Occurrence dataset https://doi.org/10.15468/ab3s5x accessed via GBIF.org on 2023-01-04 |
| download link: | https://doi.org/10.15468/dl.3x8u9c |
| *P. divaricata* ssp. laphamii (DIV) | Arizona State University Biocollections (2023). Arizona State University Vascular Plant Herbarium. Occurrence dataset https://doi.org/10.15468/a2o8vy accessed via GBIF.org on 2023-01-04. |
| | Capers R (2014). CONN. University of Connecticut. Occurrence dataset https://doi.org/10.15468/w35jmd accessed via GBIF.org on 2023-01-04. |
| | Kathryn Kalmbach Herbarium (Denver Botanic Gardens) (2022). Kathryn Kalmbach Herbarium. Occurrence dataset https://doi.org/10.15468/axrelr accessed via GBIF.org on 2023-01-04. |
| | The University of Vermont Pringle Herbarium (2023). University of Vermont, Pringle Herbarium. Occurrence dataset https://doi.org/10.15468/zsgiog accessed via GBIF.org on 2023-01-04. |
| | Franck A R, Bornhorst K (2022). University of South Florida Herbarium (USF). Version 7.373. USF Water Institute. Occurrence dataset https://doi.org/10.15468/mdnmzb accessed via GBIF.org on 2023-01-04. |
| | Grant S, von Konrat M (2022). Field Museum of Natural History (Botany) Seed Plant Collection. Version 11.14. Field Museum. Occurrence dataset https://doi.org/10.15468/nxnqzf accessed via GBIF.org on 2023-01-04. |
| | iNaturalist contributors, iNaturalist (2022). iNaturalist Research-grade Observations. iNaturalist.org. Occurrence dataset https://doi.org/10.15468/ab3s5x accessed via GBIF.org on 2023-01-04. |
| download link: | https://doi.org/10.15468/dl.2v5xt6 |
| *P. glaberrima* ssp. interior (GLA) | Kathryn Kalmbach Herbarium (Denver Botanic Gardens) (2022). Kathryn Kalmbach Herbarium. Occurrence dataset https://doi.org/10.15468/axrelr accessed via GBIF.org on 2023-01-04. |
| | Franck A R, Bornhorst K (2022). University of South Florida Herbarium (USF). Version 7.373. USF Water Institute. Occurrence dataset https://doi.org/10.15468/mdnmzb accessed via GBIF.org on 2023-01-04. |
| | The University of Vermont Pringle Herbarium (2023). University of Vermont, Pringle Herbarium. Occurrence dataset https://doi.org/10.15468/zsgiog accessed via GBIF.org on 2023-01-04. |
| | University of North Carolina at Chapel Hill Herbarium (NCU) (2023). University of North Carolina at Chapel Hill Herbarium: Vascular Plants. Occurrence dataset https://doi.org/10.15468/63vxjd accessed via GBIF.org on 2023-01-04. |
| | Roberts D (2022). CHAS Botany Collection (Arctos). Version 13.70. Chicago Academy of Sciences. Occurrence dataset https://doi.org/10.15468/ji4vbl accessed via GBIF.org on 2023-01-04. |
| | Grant S, von Konrat M (2022). Field Museum of Natural History (Botany) Seed Plant Collection. Version 11.14. Field Museum. Occurrence dataset https://doi.org/10.15468/nxnqzf accessed via GBIF.org on 2023-01-04. |
| | iNaturalist contributors, iNaturalist (2022). iNaturalist Research-grade Observations. iNaturalist.org. Occurrence dataset https://doi.org/10.15468/ab3s5x accessed via GBIF.org on 2023-01-04. |
| download link: | https://doi.org/10.15468/dl.d7ubmp |
| *P. maculata* (MAC) | E.L. Reed Herbarium at Texas Tech University (TTC) (2023). Texas Tech University, E. L. Reed Herbarium. Occurrence dataset https://doi.org/10.15468/uyakmh accessed via GBIF.org on 2023-01-04. |
| | Franck A R, Bornhorst K (2022). University of South Florida Herbarium (USF). Version 7.373. USF Water Institute. Occurrence dataset https://doi.org/10.15468/mdnmzb accessed via GBIF.org on 2023-01-04. |
| | Northern Arizona University (2023). Deaver Herbarium (Northern Arizona University). Occurrence dataset https://doi.org/10.15468/b7tfpa accessed via GBIF.org on 2023-01-04. |
| | Carnegie Museums (2023). Carnegie Museum of Natural History Herbarium. Occurrence dataset https://doi.org/10.15468/d51v1f accessed via GBIF.org on 2023-01-04. |
| | Affouard A, Joly A, Lombardo J, Champ J, Goeau H, Bonnet P (2022). Pl@ntNet automatically identified occurrences. Version 1.6. Pl@ntNet. Occurrence dataset https://doi.org/10.15468/mma2ec accessed via GBIF.org on 2023-01-04. |

| | |
|---|---|
| | Biological Collections, California State University Northridge (2023). SFV - California State University, Northridge. Occurrence dataset https://doi.org/10.15468/nrcdx7 accessed via GBIF.org on 2023-01-04. |
| | Capers R (2014). CONN. University of Connecticut. Occurrence dataset https://doi.org/10.15468/w35jmd accessed via GBIF.org on 2023-01-04. |
| | Roberts D (2022). CHAS Botany Collection (Arctos). Version 13.70. Chicago Academy of Sciences. Occurrence dataset https://doi.org/10.15468/ji4vbl accessed via GBIF.org on 2023-01-04. |
| | Grant S, von Konrat M (2022). Field Museum of Natural History (Botany) Seed Plant Collection. Version 11.14. Field Museum. Occurrence dataset https://doi.org/10.15468/nxnqzf accessed via GBIF.org on 2023-01-04. |
| | iNaturalist contributors, iNaturalist (2022). iNaturalist Research-grade Observations. iNaturalist.org. Occurrence dataset https://doi.org/10.15468/ab3s5x accessed via GBIF.org on 2023-01-04. |
| | Affouard A, Joly A, Lombardo J, Champ J, Goeau H, Bonnet P (2022). Pl@ntNet observations. Version 1.6. Pl@ntNet. Occurrence dataset https://doi.org/10.15468/gtebaa accessed via GBIF.org on 2023-01-04. |
| download link: | https://doi.org/10.15468/dl.33crrv |
| *P. paniculata* (PAN) | Orrell T, Informatics Office (2023). NMNH Extant Specimen Records (USNM, US). Version 1.65. National Museum of Natural History, Smithsonian Institution. Occurrence dataset https://doi.org/10.15468/hnhrg3 accessed via GBIF.org on 2023-01-04. |
| | Kathryn Kalmbach Herbarium (Denver Botanic Gardens) (2022). Kathryn Kalmbach Herbarium. Occurrence dataset https://doi.org/10.15468/axrelr accessed via GBIF.org on 2023-01-04. |
| | California State University, Long Beach (2023). LOB - California State University, Long Beach Herbarium. Occurrence dataset https://doi.org/10.15468/3y25yl accessed via GBIF.org on 2023-01-04. |
| | Roberts D (2022). CHAS Botany Collection (Arctos). Version 13.70. Chicago Academy of Sciences. Occurrence dataset https://doi.org/10.15468/ji4vbl accessed via GBIF.org on 2023-01-04. |
| | Carnegie Museums (2023). Carnegie Museum of Natural History Herbarium. Occurrence dataset https://doi.org/10.15468/d51v1f accessed via GBIF.org on 2023-01-04. |
| | Capers R (2014). CONN. University of Connecticut. Occurrence dataset https://doi.org/10.15468/w35jmd accessed via GBIF.org on 2023-01-04. |
| | Kerbs B, Schulenberg J (2016). H.A. Stephens Herbarium. Version 29.1. Emporia State University. Occurrence dataset https://doi.org/10.15468/k3m1qc accessed via GBIF.org on 2023-01-04. |
| | The University of Vermont Pringle Herbarium (2023). University of Vermont, Pringle Herbarium. Occurrence dataset https://doi.org/10.15468/zsgiog accessed via GBIF.org on 2023-01-04. |
| | Affouard A, Joly A, Lombardo J, Champ J, Goeau H, Bonnet P (2022). Pl@ntNet automatically identified occurrences. Version 1.6. Pl@ntNet. Occurrence dataset https://doi.org/10.15468/mma2ec accessed via GBIF.org on 2023-01-04. |
| | Northern Arizona University (2023). Deaver Herbarium (Northern Arizona University). Occurrence dataset https://doi.org/10.15468/b7tfpa accessed via GBIF.org on 2023-01-04. |
| | Grant S, von Konrat M (2022). Field Museum of Natural History (Botany) Seed Plant Collection. Version 11.14. Field Museum. Occurrence dataset https://doi.org/10.15468/nxnqzf accessed via GBIF.org on 2023-01-04. |
| | Franck A R, Bornhorst K (2022). University of South Florida Herbarium (USF). Version 7.373. USF Water Institute. Occurrence dataset https://doi.org/10.15468/mdnmzb accessed via GBIF.org on 2023-01-04. |
| | Affouard A, Joly A, Lombardo J, Champ J, Goeau H, Bonnet P (2022). Pl@ntNet observations. Version 1.6. Pl@ntNet. Occurrence dataset https://doi.org/10.15468/gtebaa accessed via GBIF.org on 2023-01-04. |
| | iNaturalist contributors, iNaturalist (2022). iNaturalist Research-grade Observations. iNaturalist.org. Occurrence dataset https://doi.org/10.15468/ab3s5x accessed via GBIF.org on 2023-01-04. |
| download link: | https://doi.org/10.15468/dl.sf37vs |
| *P. pilosa* ssp. pilosa (PIL) | Franck A R, Bornhorst K (2022). University of South Florida Herbarium (USF). Version 7.373. USF Water Institute. Occurrence dataset https://doi.org/10.15468/mdnmzb accessed via GBIF.org on 2023-01-04. |
| | Capers R (2014). CONN. University of Connecticut. Occurrence dataset https://doi.org/10.15468/w35jmd accessed via GBIF.org on 2023-01-04. |
| | iNaturalist contributors, iNaturalist (2022). iNaturalist Research-grade Observations. iNaturalist.org. Occurrence dataset https://doi.org/10.15468/ab3s5x accessed via GBIF.org on 2023-01-04. |
| | Grant S, von Konrat M (2022). Field Museum of Natural History (Botany) Seed Plant Collection. Version 11.14. Field Museum. Occurrence dataset https://doi.org/10.15468/nxnqzf accessed via GBIF.org on 2023-01-04. |
| download link: | https://doi.org/10.15468/dl.yg3nh9 |

**Table S2. Sample locations and permit information for 2022 collection**

| | | | | | |
|---|---|---|---|---|---|
| **2017-2019 collection** | | | | | |
| **Species** | **Pop.** | **Individual used** | **Lat.** | **Long.** | **Comments** |
| *Phlox divaricata* divaricata | DIV_05 | DIV-05-16 | 38.23 | -86.56 | collected by AGG [link to paper to be inserted] |
| *Phlox divaricata* divaricata | DIV_12 | DIV-12-03 | 39.87 | -86.30 | collected by AGG [link to paper to be inserted] |
| *Phlox divaricata* divaricata | DIV_31 | DIV-31-02 | 40.75 | -84.32 | collected by AGG [link to paper to be inserted] |
| *Phlox divaricata* divaricata | DIV_39 | DIV-39-1908 | 41.49 | -87.14 | collected by AGG [link to paper to be inserted] |
| *Phlox divaricata* laphamii | DIV_01 | DIV-01-02 | 37.49 | -89.35 | collected by AGG [link to paper to be inserted] |
| *Phlox divaricata* laphamii | DIV_19 | DIV-19-15 | 39.81 | -89.74 | collected by AGG [link to paper to be inserted] |
| *Phlox divaricata* laphamii | DIV_29 | DIV-29-06 | 40.22 | -88.35 | collected by AGG [link to paper to be inserted] |
| *Phlox divaricata* laphamii | DIV_39 | DIV-39-1911 | 41.49 | -87.14 | collected by AGG [link to paper to be inserted] |
| *Phlox glaberrima* interior | GLA_01 | GLA-01-02 | 40.88 | -86.88 | collected by AGG [link to paper to be inserted] |
| *Phlox glaberrima* interior | GLA_01 | GLA-01-46 | 40.88 | -86.88 | collected by AGG [link to paper to be inserted] |
| *Phlox glaberrima* interior | GLA_02 | GLA-02-11 | 41.61 | -87.55 | collected by AGG [link to paper to be inserted] |
| *Phlox glaberrima* interior | GLA_03 | GLA-03-24 | 42.00 | -87.78 | collected by AGG [link to paper to be inserted] |
| *Phlox glaberrima* interior | GLA_10 | GLA-10-1920 | 41.57 | -87.59 | collected by AGG [link to paper to be inserted] |
| *Phlox glaberrima* interior | GLA_21 | GLA-21-1921 | 40.76 | -86.66 | collected by AGG [link to paper to be inserted] |
| *Phlox maculata* | MAC_01 | MAC-01-07 | 40.35 | -87.09 | collected by AGG [link to paper to be inserted] |

| | | | | | | | | | | | | | |
|---|---|---|---|---|---|---|---|---|---|---|---|---|---|
| *Phlox maculata* | MAC_01 | | MAC-01-1910 | 40.35 | -87.09 | | | | | | | | collected by AGG [link to paper to be inserted] |
| *Phlox maculata* | MAC_02 | | MAC-02-1902 | 39.28 | -86.46 | | | | | | | | collected by AGG [link to paper to be inserted] |
| *Phlox maculata* | MAC_02 | | MAC-02-1910 | 39.28 | -86.46 | | | | | | | | collected by AGG [link to paper to be inserted] |
| *Phlox paniculata* | PAN_01 | | PAN-01-1901 | 40.35 | -87.09 | | | | | | | | collected by AGG [link to paper to be inserted] |
| *Phlox paniculata* | PAN_02 | | PAN-02-1917 | 40.67 | -86.63 | | | | | | | | collected by AGG [link to paper to be inserted] |
| *Phlox paniculata* | PAN_03 | | PAN-03-08 | 39.16 | -86.31 | | | | | | | | collected by AGG [link to paper to be inserted] |
| *Phlox paniculata* | PAN_07 | | PAN-07-1906 | 41.57 | -87.59 | | | | | | | | collected by AGG [link to paper to be inserted] |
| *Phlox pilosa* WHITE | PIL_20 | | PIL-20-1922 | 41.57 | -87.60 | | | | | | | | collected by AGG [link to paper to be inserted] |
| *Phlox pilosa* pilosa | PIL_06 | | PIL-06-1951 | 40.45 | -88.10 | | | | | | | | collected by AGG [link to paper to be inserted] |
| *Phlox pilosa* pilosa | PIL_09 | | PIL-09-01 | 40.67 | -87.39 | | | | | | | | collected by AGG [link to paper to be inserted] |
| *Phlox pilosa* pilosa | PIL_10 | | PIL-10-92 | 41.17 | -87.11 | | | | | | | | collected by AGG [link to paper to be inserted] |
| *Phlox pilosa* pilosa | PIL_21 | | PIL-21-05 | 41.60 | -87.45 | | | | | | | | collected by AGG [link to paper to be inserted] |
| *Phlox pilosa* sangamonensis | SAN_01 | | SAN-01-07 | 40.23 | -88.35 | | | | | | | | collected by AGG [link to paper to be inserted] |
| *Phlox pilosa* sangamonensis | SAN_01 | | SAN-01-17 | 40.23 | -88.35 | | | | | | | | collected by AGG [link to paper to be inserted] |
| *Phlox pilosa* sangamonensis | SAN_03 | | SAN-03-12 | 40.21 | -88.38 | | | | | | | | collected by AGG [link to paper to be inserted] |
| **2022 collection** | | | | | | | | | | | | | |
| **Species** | **Pop. code AFF** | **Original pop. code AGG** | **Individual used** | **Lat.** | **Long.** | **# Samples collected** | **Date** | **State** | **Forest/ park/ preserve** | **County/ district/ department** | **Permit number** | **Local permit by** | **Comments** |
| *Phlox divaricata* divaricata | DIV-1-AFF | DIV-17? | DIV-1-7-aff | 39.890 | -87.197 | 16 | 0512 | IN | Turkey Run State Park | Indiana Department of Natural Resources | NP22-48 | - | the permit by Indiana Division of Nature Preserves includes Nature Preserve section of the listed parks |
| *Phlox divaricata* divaricata | DIV-2-AFF | DIV-12 | | 39.868 | -86.297 | 21 | 0513 | IN | Eagle Creek Park | Indiana Department of Natural Resources | NP22-48 | Eagle Creek Park | the permit by Indiana Division of Nature Preserves includes Nature Preserve section of the listed parks |

| | | | | | | | | | | | | | |
|---|---|---|---|---|---|---|---|---|---|---|---|---|---|
| *Phlox divaricata* divaricata | DIV-3-AFF | DIV-11? | | 39.204 | 86.346 | 8 | 0514 | IN | Yellowwod State Forest | Indiana Department of Natural Resources | NP22-48 | - | the permit by Indiana Division of Nature Preserves includes Nature Preserve section of the listed parks |
| *Phlox divaricata* divaricata | DIV-4-AFF | DIV-05 | | 38.230 | 86.560 | 6 | 0515 | IN | roadside (in foresty patch) | - | - | - | |
| *Phlox divaricata* divaricata | DIV-5-AFF | DIV-31 | | 40.749 | -84.324 | 18 | 0524 | OH | Kendrick Woods | Johnny Appleseed Metropolitan Park District | - | - | |
| *Phlox divaricata* laphamii | LAP-1-AFF | DIV-01 | | 37.490 | -89.350 | 2 | 0516 | IL | Giant City State Park | Illinois Department of Natural Resources | # SS22-054 | - | |
| *Phlox divaricata* laphamii | LAP-2-AFF | DIV-01 | | 37.490 | -89.350 | 2 | 0516 | IL | Trail of Tears State Forest | Illinois Department of Natural Resources | # SS22-054 | - | |
| *Phlox divaricata* laphamii | LAP-3-AFF | DIV-01 | | 37.490 | -89.350 | 3 | 0516 | IL | Trail of Tears State Forest | Illinois Department of Natural Resources | # SS22-054 | - | |
| *Phlox divaricata* laphamii | LAP-4-AFF | DIV-01 | | 37.490 | -89.350 | 5 | 0516 | IL | Trail of Tears State Forest | Illinois Department of Natural Resources | # SS22-054 | - | |
| *Phlox divaricata* laphamii | LAP-5-AFF | DIV-18 or DIV 30? | | 40.206 | -88.384 | 16 | 0519 | IL | Lake of the Woods Forest Preserve | Champaign County Forest Preserve District | - | - | |
| *Phlox divaricata* laphamii | LAP-6-AFF | N/A | | 40.139 | -88.472 | 9 | 0519 | IL | roadside (in foresty patch near Sangamon park) | - | - | - | |
| *Phlox divaricata* laphamii | LAP-7-AFF | N/A | | 41.570 | -87.593 | 10 | 0521 | IL | Wampun Lake / Thornton Lansing Road Nature Preserve | Illinois Department of Natural Resources | # SS22-054 | Forest Preserves of Cook County | plus Nature Preserves Permit, issued by Illinois Nature Preserves Commission |
| *Phlox glaberrima* interior | GLA-1-AFF | GLA-10 | | 41.568 | -87.593 | 15 | 0703 | IL | Thornton Lansing Road Nature Preserve | Illinois Department of Natural Resources | # SS22-054 | Forest Preserves of Cook County | |
| *Phlox glaberrima* interior | GLA-2-AFF | GLA-02 | GLA-2-2-aff | 41.614 | -87.550 | 10 | 0703 | IL | Sand Ridge Nature Preserve | Illinois Department of Natural Resources | # SS22-054 | Forest Preserves of Cook County | |
| *Phlox glaberrima* interior | GLA-3-AFF | GLA-03 | | 42.002 | 87.779 | 20 | 0703 | IL | Sydney Yates Flatwoods | Illinois Department of Natural Resources | # SS22-054 | Forest Preserves of Cook County | |

| | | | | | | | | | | | | | |
|---|---|---|---|---|---|---|---|---|---|---|---|---|---|
| *Phlox maculata* | MAC-1-AFF | MAC-01 | | 40.354 | -87.091 | 18 | 0701 | IN | roadside | - | - | - | |
| *Phlox paniculata* | PAN-1-AFF | N/A | | 39.281 | -86.463 | 1 | 0630 | IN | roadside | - | - | - | |
| *Phlox paniculata* | PAN-2-AFF | PAN-04 | | 39.890 | -87.199 | 3 | 0701 | IN | Turkey Run State Park | Indiana Department of Natural Resources | NP22-48 | - | the permit by Indiana Division of Nature Preserves includes Nature Preserve section of the listed parks |
| *Phlox paniculata* | PAN-3-AFF | PAN-01 | PAN-3-3-aff | 40.354 | -87.091 | 3 | 0701 | IN | roadside | - | - | - | |
| *Phlox paniculata* | PAN-4-AFF | PAN-02 | PAN-4-3-aff | 40.673 | -86.630 | 12 | 0702 | IN | roadside | - | - | - | |
| *Phlox pilosa* pilosa | PIL-1-AFF | PIL-06 | | 40.446 | -88.098 | 20 | 0520 | IL | Prospect Cemetery Nature Preserve | Illinois Department of Natural Resources | # SS22-054 | Grand Prarie Friends | plus Nature Preserves Permit, issued by Illinois Nature Preserves Commission |
| *Phlox pilosa* pilosa | PIL-2-AFF | PIL-09 | | 40.670 | -87.390 | 14 | 0520 | IN | roadside (between gravel road and railroad) | - | - | - | |
| *Phlox pilosa* pilosa | PIL-3-AFF | PIL-21 | | 41.600 | -87.450 | 4 | 0521 | IN | Gibson Woods Nature Preserve | Indiana Department of Natural Resources | NP22-48 | - | the permit by Indiana Division of Nature Preserves includes Nature Preserve section of the listed parks |
| *Phlox pilosa* pilosa | PIL-4-AFF | PIL-10 | | 41.173 | -87.102 | 19 | 0522 | IN | Stoutsburg Savanna Nature Preserve | Indiana Department of Natural Resources | NP22-48 | - | the permit by Indiana Division of Nature Preserves includes Nature Preserve section of the listed parks |

**Table S3. Crosses overview**

| Female individual ID | Female species abbr. | Male individual ID | Male species abbr. | Cross name | Cross abbr. | Cross type | Crossed flowers |
|---|---|---|---|---|---|---|---|
| LAP-DIV-19-15 | LAP | DIV-05-16 | DIV | LAP-DIV-19-15_DIV-05-16 | DIVxDIV | het | 10 |
| DIV-31-02 | DIV | DIV-05-16 | DIV | DIV-31-02_DIV-05-16 | DIVxDIV | con | 9 |
| LAP-DIV-01-02 | LAP | LAP-DIV-19-15 | LAP | LAP-DIV-01-02_LAP-DIV-19-15 | DIVxDIV | con | 7 |
| DIV-31-02 | DIV | GLA-21-1921 | GLA | DIV-31-02_GLA-21-1921 | DIVxGLA | het | 10 |
| LAP-DIV-19-15 | LAP | GLA-02-11 | GLA | LAP-DIV-19-15_GLA-02-11 | DIVxGLA | het | 10 |
| LAP-DIV-29-06 | LAP | GLA-01-02 | GLA | LAP-DIV-29-06_GLA-01-02 | DIVxGLA | het | 10 |
| LAP-DIV-19-15 | LAP | MAC-02-1910 | MAC | LAP-DIV-19-15_MAC-02-1910 | DIVxMAC | het | 10 |
| LAP-DIV-01-02 | LAP | MAC-01-07 | MAC | LAP-DIV-01-02_MAC-01-07 | DIVxMAC | het | 10 |
| LAP-DIV-19-15 | LAP | MAC-02-1902 | MAC | LAP-DIV-19-15_MAC-02-1902 | DIVxMAC | het | 7 |
| LAP-DIV-19-15 | LAP | PAN-01-1901 | PAN | LAP-DIV-19-15_PAN-01-1901 | DIVxPAN | het | 10 |
| DIV-12-03 | DIV | PAN-07-1906 | PAN | DIV-12-03_PAN-07-1906 | DIVxPAN | het | 10 |
| LAP-DIV-19-15 | LAP | PAN-03-08 | PAN | LAP-DIV-19-15_PAN-03-08 | DIVxPAN | het | 10 |
| DIV-31-02 | DIV | PIL-06-1951 | PIL | DIV-31-02_PIL-06-1951 | DIVxPIL | het | 10 |
| LAP-DIV-39-1911 | LAP | PIL-20-1922 | PIL | LAP-DIV-39-1911_PIL-20-1922 | DIVxPIL | het | 6 |
| LAP-DIV-19-15 | LAP | PIL-10-92 | PIL | LAP-DIV-19-15_PIL-10-92 | DIVxPIL | het | 10 |
| GLA-03-24 | GLA | DIV-31-02 | DIV | GLA-03-24_DIV-31-02 | GLAxDIV | het | 10 |
| GLA-10-1920 | GLA | DIV-05-16 | DIV | GLA-10-1920_DIV-05-16 | GLAxDIV | het | 10 |
| GLA-03-24 | GLA | LAP-DIV-01-02 | LAP | GLA-03-24_LAP-DIV-01-02 | GLAxDIV | het | 10 |
| GLA-03-24 | GLA | GLA-02-11 | GLA | GLA-03-24_GLA-02-11 | GLAxGLA | con | 10 |
| GLA-21-1921 | GLA | GLA-03-24 | GLA | GLA-21-1921_GLA-03-24 | GLAxGLA | con | 10 |
| GLA-01-02 | GLA | GLA-10-1920 | GLA | GLA-01-02_GLA-10-1920 | GLAxGLA | con | 10 |
| GLA-10-1920 | GLA | MAC-02-1902 | MAC | GLA-10-1920_MAC-02-1902 | GLAxMAC | het | 10 |
| GLA-02-11 | GLA | MAC-01-07 | MAC | GLA-02-11_MAC-01-07 | GLAxMAC | het | 10 |
| GLA-01-02 | GLA | MAC-02-1910 | MAC | GLA-01-02_MAC-02-1910 | GLAxMAC | het | 10 |
| GLA-01-02 | GLA | PAN-01-1901 | PAN | GLA-01-02_PAN-01-1901 | GLAxPAN | het | 10 |
| GLA-10-1920 | GLA | PAN-07-1906 | PAN | GLA-10-1920_PAN-07-1906 | GLAxPAN | het | 10 |
| GLA-21-1921 | GLA | PAN-03-08 | PAN | GLA-21-1921_PAN-03-08 | GLAxPAN | het | 10 |
| GLA-02-11 | GLA | PIL-06-1951 | PIL | GLA-02-11_PIL-06-1951 | GLAxPIL | het | 10 |
| GLA-10-1920 | GLA | PIL-20-1922 | PIL | GLA-10-1920_PIL-20-1922 | GLAxPIL | het | 10 |
| GLA-21-1921 | GLA | PIL-10-92 | PIL | GLA-21-1921_PIL-10-92 | GLAxPIL | het | 10 |
| MAC-02-1902 | MAC | LAP-DIV-01-02 | LAP | MAC-02-1902_LAP-DIV-01-02 | MACxDIV | het | 10 |
| MAC-02-1910 | MAC | DIV-12-03 | DIV | MAC-02-1910_DIV-12-03 | MACxDIV | het | 10 |
| MAC-02-1910 | MAC | LAP-DIV-19-15 | LAP | MAC-02-1910_LAP-DIV-19-15 | MACxDIV | het | 10 |
| MAC-02-1902 | MAC | GLA-02-11 | GLA | MAC-02-1902_GLA-02-11 | MACxGLA | het | 10 |
| MAC-01-07 | MAC | GLA-01-02 | GLA | MAC-01-07_GLA-01-02 | MACxGLA | het | 10 |
| MAC-01-1910 | MAC | GLA-21-1921 | GLA | MAC-01-1910_GLA-21-1921 | MACxGLA | het | 10 |
| MAC-02-1910 | MAC | MAC-01-07 | MAC | MAC-02-1910_MAC-01-07 | MACxMAC | con | 10 |
| MAC-02-1902 | MAC | MAC-01-07 | MAC | MAC-02-1902_MAC-01-07 | MACxMAC | con | 10 |
| MAC-01-07 | MAC | MAC-02-1902 | MAC | MAC-01-07_MAC-02-1902 | MACxMAC | con | 10 |
| MAC-02-1902 | MAC | PAN-03-08 | PAN | MAC-02-1902_PAN-03-08 | MACxPAN | het | 10 |
| MAC-01-1910 | MAC | PAN-01-1901 | PAN | MAC-01-1910_PAN-01-1901 | MACxPAN | het | 10 |
| MAC-01-07 | MAC | PAN-07-1906 | PAN | MAC-01-07_PAN-07-1906 | MACxPAN | het | 10 |
| MAC-02-1902 | MAC | PIL-20-1922 | PIL | MAC-02-1902_PIL-20-1922 | MACxPIL | het | 10 |
| MAC-01-1910 | MAC | PIL-10-92 | PIL | MAC-01-1910_PIL-10-92 | MACxPIL | het | 10 |
| MAC-01-07 | MAC | PIL-21-05 | PIL | MAC-01-07_PIL-21-05 | MACxPIL | het | 10 |
| PAN-01-1901 | PAN | LAP-DIV-19-15 | LAP | PAN-01-1901_LAP-DIV-19-15 | PANxDIV | het | 10 |
| PAN-07-1906 | PAN | LAP-DIV-39-1911 | LAP | PAN-07-1906_LAP-DIV-39-1911 | PANxDIV | het | 7 |
| PAN-03-08 | PAN | DIV-12-03 | DIV | PAN-03-08_DIV-12-03 | PANxDIV | het | 4 |
| PAN-01-1901 | PAN | GLA-21-1921 | GLA | PAN-01-1901_GLA-21-1921 | PANxGLA | het | 10 |
| PAN-03-08 | PAN | GLA-10-1920 | GLA | PAN-03-08_GLA-10-1920 | PANxGLA | het | 10 |
| PAN-07-1906 | PAN | GLA-02-11 | GLA | PAN-07-1906_GLA-02-11 | PANxGLA | het | 10 |
| PAN-01-1901 | PAN | MAC-01-07 | MAC | PAN-01-1901_MAC-01-07 | PANxMAC | het | 10 |
| PAN-03-08 | PAN | MAC-02-1902 | MAC | PAN-03-08_MAC-02-1902 | PANxMAC | het | 10 |
| PAN-07-1906 | PAN | MAC-02-1910 | MAC | PAN-07-1906_MAC-02-1910 | PANxMAC | het | 10 |
| PAN-03-08 | PAN | PAN-01-1901 | PAN | PAN-03-08_PAN-01-1901 | PANxPAN | con | 10 |
| PAN-01-1901 | PAN | PAN-07-1906 | PAN | PAN-01-1901_PAN-07-1906 | PANxPAN | con | 10 |
| PAN-07-1906 | PAN | PAN-03-08 | PAN | PAN-07-1906_PAN-03-08 | PANxPAN | con | 10 |
| PAN-01-1901 | PAN | PIL-20-1922 | PIL | PAN-01-1901_PIL-20-1922 | PANxPIL | het | 10 |
| PAN-01-1901 | PAN | PIL-10-92 | PIL | PAN-01-1901_PIL-10-92 | PANxPIL | het | 10 |

| | | | | | | | |
|---|---|---|---|---|---|---|---|
| PAN-02-1917 | PAN | PIL-09-01 | PIL | PAN-02-1917_PIL-09-01 | PANxPIL | het | 10 |
| PIL-06-1951 | PIL | DIV-31-02 | DIV | PIL-06-1951_DIV-31-02 | PILxDIV | het | 10 |
| PIL-10-92 | PIL | LAP-DIV-39-1911 | LAP | PIL-10-92_LAP-DIV-39-1911 | PILxDIV | het | 7 |
| PIL-21-05 | PIL | LAP-DIV-01-02 | LAP | PIL-21-05_LAP-DIV-01-02 | PILxDIV | het | 4 |
| PIL-06-1951 | PIL | GLA-02-11 | GLA | PIL-06-1951_GLA-02-11 | PILxGLA | het | 10 |
| PIL-21-05 | PIL | GLA-03-24 | GLA | PIL-21-05_GLA-03-24 | PILxGLA | het | 10 |
| PIL-20-1922 | PIL | GLA-21-1921 | GLA | PIL-20-1922_GLA-21-1921 | PILxGLA | het | 10 |
| PIL-10-92 | PIL | MAC-02-1902 | MAC | PIL-10-92_MAC-02-1902 | PILxMAC | het | 10 |
| PIL-20-1922 | PIL | MAC-01-1910 | MAC | PIL-20-1922_MAC-01-1910 | PILxMAC | het | 10 |
| PIL-21-05 | PIL | MAC-01-07 | MAC | PIL-21-05_MAC-01-07 | PILxMAC | het | 10 |
| PIL-20-1922 | PIL | PAN-07-1906 | PAN | PIL-20-1922_PAN-07-1906 | PILxPAN | het | 10 |
| PIL-10-92 | PIL | PAN-03-08 | PAN | PIL-10-92_PAN-03-08 | PILxPAN | het | 10 |
| PIL-06-1951 | PIL | PAN-01-1901 | PAN | PIL-06-1951_PAN-01-1901 | PILxPAN | het | 5 |
| PIL-20-1922 | PIL | PIL-21-05 | PIL | PIL-20-1922_PIL-21-05 | PILxPIL | con | 10 |
| PIL-06-1951 | PIL | PIL-20-1922 | PIL | PIL-06-1951_PIL-20-1922 | PILxPIL | con | 10 |
| PIL-21-05 | PIL | PIL-06-1951 | PIL | PIL-21-05_PIL-06-1951 | PILxPIL | con | 10 |

**Table S4. Cross results and RI values.** Table is sorted from lowest RI values to highest and alphabetically by cross where the values are the same.

| **Cross** | **Cross type** | **Average number of seeds per flower** | **RI** |
|---|---|---|---|
| DIVxDIV | conspecific | 1.52 | n/a |
| PILxPIL | conspecific | 1.40 | n/a |
| GLAxGLA | conspecific | 1.43 | n/a |
| MACxMAC | conspecific | 1.37 | n/a |
| PANxPAN | conspecific | 0.97 | n/a |
| MACxGLA | long x long | 0.80 | 0.262 |
| GLAxMAC | long x long | 0.67 | 0.365 |
| DIVxGLA | short x long | 0.70 | 0.368 |
| PILxGLA | short x long | 0.63 | 0.377 |
| PANxGLA | long x long | 0.40 | 0.415 |
| DIVxPIL | short x short | 0.30 | 0.670 |
| GLAxPAN | long x long | 0.27 | 0.686 |
| PILxPAN | short x long | 0.20 | 0.750 |
| PANxMAC | long x long | 0.07 | 0.871 |
| DIVxPAN | short x long | 0.10 | 0.876 |
| PILxDIV | short x short | 0.07 | 0.909 |
| DIVxMAC | short x long | 0.07 | 0.916 |
| GLAxDIV | long x short | 0.00 | 1.000 |
| GLAxPIL | long x short | 0.00 | 1.000 |
| MACxDIV | long x short | 0.00 | 1.000 |
| MACxPAN | long x long | 0.00 | 1.000 |
| MACxPIL | long x short | 0.00 | 1.000 |
| PANxDIV | long x short | 0.00 | 1.000 |
| PANxPIL | long x short | 0.00 | 1.000 |
| PILxMAC | short x long | 0.00 | 1.000 |
